# HLA class I genotypes customize vaccination strategies in immune simulation to combat COVID-19

**DOI:** 10.1101/2020.11.18.388983

**Authors:** Shouxiong Huang, Ming Tan

## Abstract

Memory CD8^+^ T cells are associated with a better outcome in Coronavirus Disease 2019 (COVID-19) and recognized as promising vaccine targets against viral infections. This study determined the efficacy of population-dominant and infection-relevant human leukocyte antigens (HLA) class I proteins to present severe acute respiratory syndrome coronavirus 2 (SARS-CoV-2) peptides through calculating binding affinities and simulating CD8^+^ T cell responses. As a result, HLA class I proteins distinguished or shared various viral peptides derived from viruses. HLA class I supertypes clustered viral peptides through recognizing anchor and preferred residues. SARS-CoV-2 peptides overlapped significantly with SARS but minimally with common human coronaviruses. Immune simulation of CD8^+^ T cell activation using predicted SARS-CoV-2 peptide antigens depended on high-affinity peptide binding, anchor residue interaction, and synergistic presentation of HLA class I proteins in individuals. Results demonstrated that multi-epitope vaccination, employing a strong binding affinity, viral adjuvants, and heterozygous HLA class I genes, induced potent immune responses. Therefore, optimal CD8^+^ T cell responses can be achieved and customized contingent on HLA class I genotypes in human populations, supporting a precise vaccination strategy to combat COVID-19.

## Introduction

Coronavirus disease 2019 (COVID-19) pandemics has raised more than 55 million cases globally since earlier this year and remains rapid spreading with over a half-million new daily infections worldwide^1^. COVID-19 is caused by the infection of severe acute respiratory syndrome coronavirus 2 (SARS-CoV-2), characterized as pneumonia and lymphopenia, and develops severe respiratory or multiple organ failure in around 5% of patients^2^. Recent vaccine development mainly focused on the induction of neutralizing antibodies. Although multiple vaccination strategies are under clinical trials^3^, their protection against COVID-19 in large human populations remains unknown. Considering disappointing efficacy or insufficient protection frequently occurred in previous anti-viral vaccine trials using large human populations, alternative approaches to stimulate protective immunity and control infections effectively remain urgently needed.

CD8^+^ T cells mediate essential immune responses against numerous intracellular viral infections to provide durable T cell memory and protection up to many years after infections, such as in immune defense against SARS^4–6^, HIV^7,8^, and influenza virus infections^9,10^. Clinical trials of anti-viral vaccines frequently led to disappointing results with weak CD8^+^ T cell responses^7,8,11^, supporting the importance of anti-viral CD8^+^ T cell responses in different human populations. Indeed, viral-specific CD8^+^ T cells are essential for viral clearance and long-term protection against HIV^12^, Ebola^13^, hepatitis B viruses^14^, and cytomegalovirus^15^, despite the critical role of neutralizing antibodies for blocking viral entry to human cells^6,16–18^. In COVID-19, it is likely important to stimulate both neutralizing antibodies and CD8^+^ T cell responses to achieve full protection, as in viral vaccination trials against SARS^19^ and influenza virus^6,20^. SARS-CoV-2 infections induce a dramatic reduction of CD8^+^ T cells associated with severe^21,22^ and elderly patients^23^. Multiple recent reports strongly support that viral-specific and memory CD8^+^ T cell responses are associated with milder diseases, younger patients^24^, and convalescent phases of infections^25,26^, although HLA genotype^27^, differentiation kinetics^28,29^, and adjuvant usage^30^ impact the protection of CD8^+^ T cells subsets.

Activation of viral-specific CD8^+^ T cells and killing of viral-infected cells rely on antigen-presentation mediated by HLA class I proteins together with co-stimulation and cytokine microenvironment^31,32^. A fundamental mechanism is that HLA class I proteins bind specific viral peptides as in a key-to-lock model^33,34^, which interacts with the T cell receptor of CD8^+^ T cells for activation and viral-infected cell recognition. Similar to the cell entry of SARS-CoV^35^, SARS-CoV-2 is considered to enter lung epithelial cells and blood cells through interacting with cell surface receptors, such as angiotensin-converting enzyme 2 (ACE2), and follow the endocytic and cytoplasmic compartments^36^. Intracellular viral proteins in viral assembling and trafficking processes can be degraded in proteasome or endosome to form antigenic peptide ligands that further bind to HLA class I proteins through a secretory pathway or cross-presentation^37,38^. Currently, computational prediction and experimental identification of peptide antigens for CD8^+^ T cell activation in COVID-19 are ongoing for a purpose of “identifying vaccine candidates”^26,39–41^. However, the impact of HLA class I genotypes on peptide antigen selection and CD8^+^ T cell activation remains poorly understood.

To date, the HLA-A locus displays 6291 alleles that encode around 3900 proteins, and HLA-B displays 7562 alleles that encode around 4808 proteins in humans^42^. It is unlikely to fully unveil functional CD8^+^ T cell responses to unique peptide sets presented by each HLA class I protein. With a wealth of structural knowledge on HLA-peptide-T cell receptor interaction^32,43^, computational prediction and immune simulation as a prototypic artificial intelligent tool become possible and essential to facilitate structural identification and functional simulation of peptide antigen presentation for CD8^+^ T cell activation. This study applied affinity binding theory to predict SARS-CoV-2 peptides for binding to different HLA class I proteins and simulate the antigen-presentation process for CD8^+^ T cell activation. Results showed a dramatically different capacity in various HLA class I proteins to bind SARS-CoV-2 peptides. Multiple HLA class I proteins that have been indicated to associate with more severe symptoms or weaker CD8^+^ T cell responses in SARS^44^ were demonstrated in this study for binding a less number of SARS-CoV-2 peptides. In contrast, HLA class I proteins that have been suggested protective or potent stimulators of CD8^+^ T cells in SARS^45^ were predicted to bind a large number of SARS-CoV-2 peptides. Different HLA class I proteins clustered to a supertype share identical sequences and similar binding patterns of SARS-CoV-2 peptides. Across various coronaviruses, a large number of peptides from SARS viruses and a limited number of peptides from “common cold”-causing human coronaviruses are identical to those of SARS-CoV-2. Further, affinity-based immune simulation provided an optimized combination of peptides dependent on HLA class I genotypes to facilitate customizing precise vaccination strategies for individuals and human populations.

## Results

### Dominant HLA class I proteins sample various numbers of SARS-CoV-2 peptides

To stimulate anti-COVID-19 CD8^+^ T cell responses, this study used NetMHCpan program to comprehensively determine the binding affinity of SARS-CoV-2 peptides to ten dominant HLA class I A and class I B proteins, which are frequently detected in the United States (US)^46^. HLA-C alleles were not involved since HLA-C proteins mainly induce immune tolerance^47^. The threshold of sufficient binding affinity was defined using a half-maximal inhibitory concentration (IC50<500nM), in which the maximal concentration of a SARS-CoV-2 peptide is required to competing off the binding of a probe peptide to each HLA class I protein. The spike (S), membrane (M), nucleocapsid (N), and multiple ORF-encoded proteins (Fig. S1A) of the SARS-CoV-2 virus are considered immunogenic and are analyzed in this study because representative peptides of these proteins stimulate CD8^+^ T cells isolated from COVID-19 patients^39,40^. Various numbers of SARS-CoV-2 peptides were predicted with a high binding affinity to ten dominant HLA class I A and B proteins (Fig. S1A). These ten alleles exist in 78.5% of the US population approximately. Upon the further consideration of the earlier processes in antigen presentation, including proteasome degradation (cutoff>5 potential cutting sites by proteasome enzymes)^48^ and peptide transportation through the transporter associated with antigen processing (TAP, cutoff IC50<1nM)^49^, 80% to 100% peptide ligands for HLA class I proteins remained significant (Fig. 1A).

**Fig. 1.**
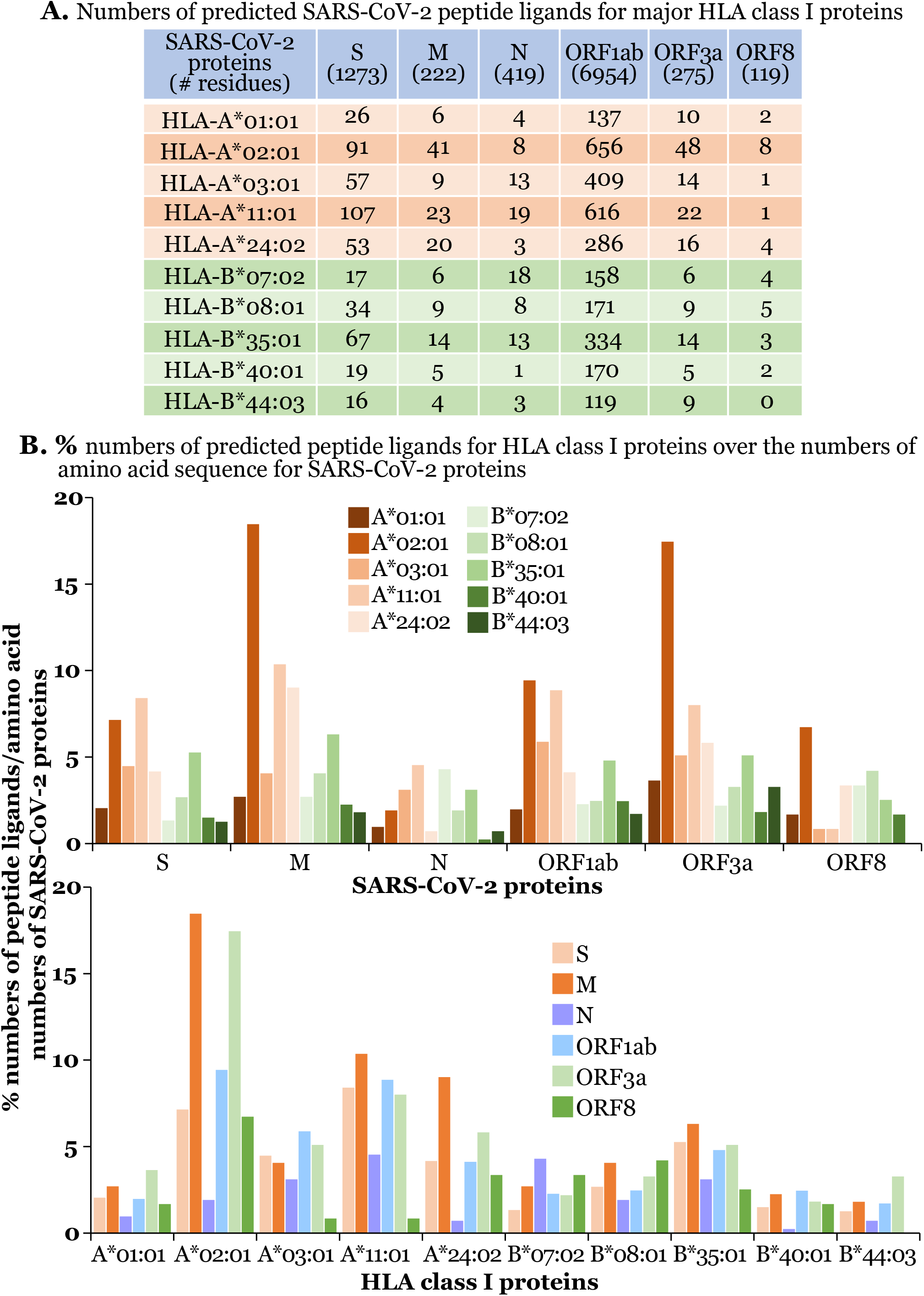
Dominant HLA class I proteins sampled various numbers of SARS-CoV-2 peptides. Peptide ligands were predicted based on binding affinity to HLA class I proteins, digestion sites for proteasome enzymes, and binding affinity to TAP transporter. The numbers of predicted peptide ligands from immunogenic SARS-CoV-2 proteins are listed for ten dominant HLA class I proteins **(A)**. % of the numbers of predicted peptides over the amino acid numbers of different SARS-CoV-2 proteins are plotted to reflect the relative frequencies of HLA class I peptide ligands **(B)**.

As a result, striking variation exists in the numbers of SARS-CoV-2 peptides sampled by different dominant HLA class I proteins. Most of the ten dominant HLA-A and B proteins bound high numbers of peptides derived from noted immunogenic SARS-CoV-2 proteins. A*02:01 and A*11:01 bound the highest numbers of peptides (Fig. 1), indicating that a large human population expressing at least one of these dominant HLA class I alleles is able to respond to SARS-CoV-2 infection. The capacity of binding a higher number of viral peptides suggests a potential dominancy of viral-induced CD8^+^ T cell responses. These potent peptide binders can compensate with weak peptide binders and balance the overall ability of antigen presentation for CD8^+^ T cell activation, if an HLA class I genotype combines both potent and weak peptide binders in the same individual. Apparently, some SARS-CoV-2 proteins provided a lower number of peptides due to their short chain length of proteins, such as N and ORF8-encoded proteins. Thus, to estimate a critical protein source of viral peptides, peptide numbers were further normalized with the chain length of each SARS-CoV-2 protein (Fig. 1B). As a result, among different SARS-CoV-2 proteins, fewer percentage of peptides from N and ORF8-encoded proteins are presented by most HLA-A proteins. Among different HLA class I proteins, HLA-A proteins overall displayed a higher peptide loading capacity than HLA-B proteins. For example, HLA-A*02:01, A*11:01, and A*24:02 sampled a higher % of peptides from multiple viral proteins, except for N and ORF8-encoded proteins (Fig. 1B). HLA-B*35:01 showed a high peptide loading capacity than other HLA-B proteins. Although the numbers of peptide ligands do not necessarily reflect the protection of peptide antigens in diseases, predicted peptides binding to HLA proteins raise a critical concern that various human populations with various HLA class I genotypes will respond differently in COVID-19 infections.

To consider whether HLA class I sequence heterogeneity selects various numbers of peptide ligands, α1 and α2 ligand-binding domains of ten HLA class I proteins were aligned (Fig. S1B), using a hierarchical clustering approach^50^. Overall resulted similarity ranged from 75% to 96%, with the highest similarity between A*03:01 and A*11:01 (96.15%) (Fig. S1C). However, the numbers of predicted peptides appear generally unmatched with the degree of similarity between different HLA class I proteins. For example, A*02:01 shows 92.31% similarity to A*03:01, but they have very different numbers of binding peptides for the same proteins (Fig. 1A). Similar non-concordance occurs between A*01:01 and A*11:01 with 93.96% sequence similarity and between A*03:01 and A*11:01 with the highest sequence similarity of 96.15%. These results suggest a need to exam the similarity of peptide-binding domains among HLA class I proteins.

### Dominant HLA class I proteins share limited numbers of SARS-CoV-2 peptides

To determine the conservation of viral peptides for CD8^+^ T cell activation in different human populations, we applied Venn diagrams to show the numbers of predicted identical peptides between dominant HLA-A proteins (Fig. 2A) and HLA-B proteins (Fig. 2B). We used the spike and ORF1ab genes, which encode longer protein sequences than others, to provide a higher chance of overlapped peptide sampling by different HLA class I proteins. As a result, identical peptides are infrequently shared by many HLA class I proteins despite high sequence similarity (>90%) between HLA-A or HLA-B proteins (Fig. S1C). For some HLA proteins, vast numbers of shared peptides were observed, for example, between A*03:01 and A*11:01 with an extremely high sequence similarity of 96.15%, across B*40:01 and B*44:03 with a relatively low similarity of 89.56%, or between B*35:01 and B*07:02 with a lower sequence similarity of 88.46%. In particular, almost 80% or 85% of peptides bound to A*03:01 also bind to A*11:01. However, HLA sequence similarity is overall discordant to the numbers of shared peptides. B*08:01 and B*07:02 (93.41% sequence similarity) shared much fewer numbers of peptides, and B*08:01 and B*40:01 (92.31% sequence similarity) shared no peptide from the S protein or only 1 out of 340 peptides from the ORF1ab-encoded proteins. Therefore, an overall high sequence similarity does not suggest a high number of peptide overlap. As in the sequence alignment (Fig. S1B), potential determinants for the numbers of shared peptides likely originate from hot spots with limited numbers of residues in ligand-binding grooves. The similarity of these key residues and pocket structures assigns HLA class I proteins to multiple supertypes^51,52^. Indeed, high numbers of identical SARS-CoV-2 peptides were shared between HLA class I proteins of the same supertypes, such as supertypes A03 (A*03:01 and A*11:01), B07 (B*35:01 and B*07:02), and B40 (B*40:01 and B*44:03) (Fig. 2).

**Fig. 2.**
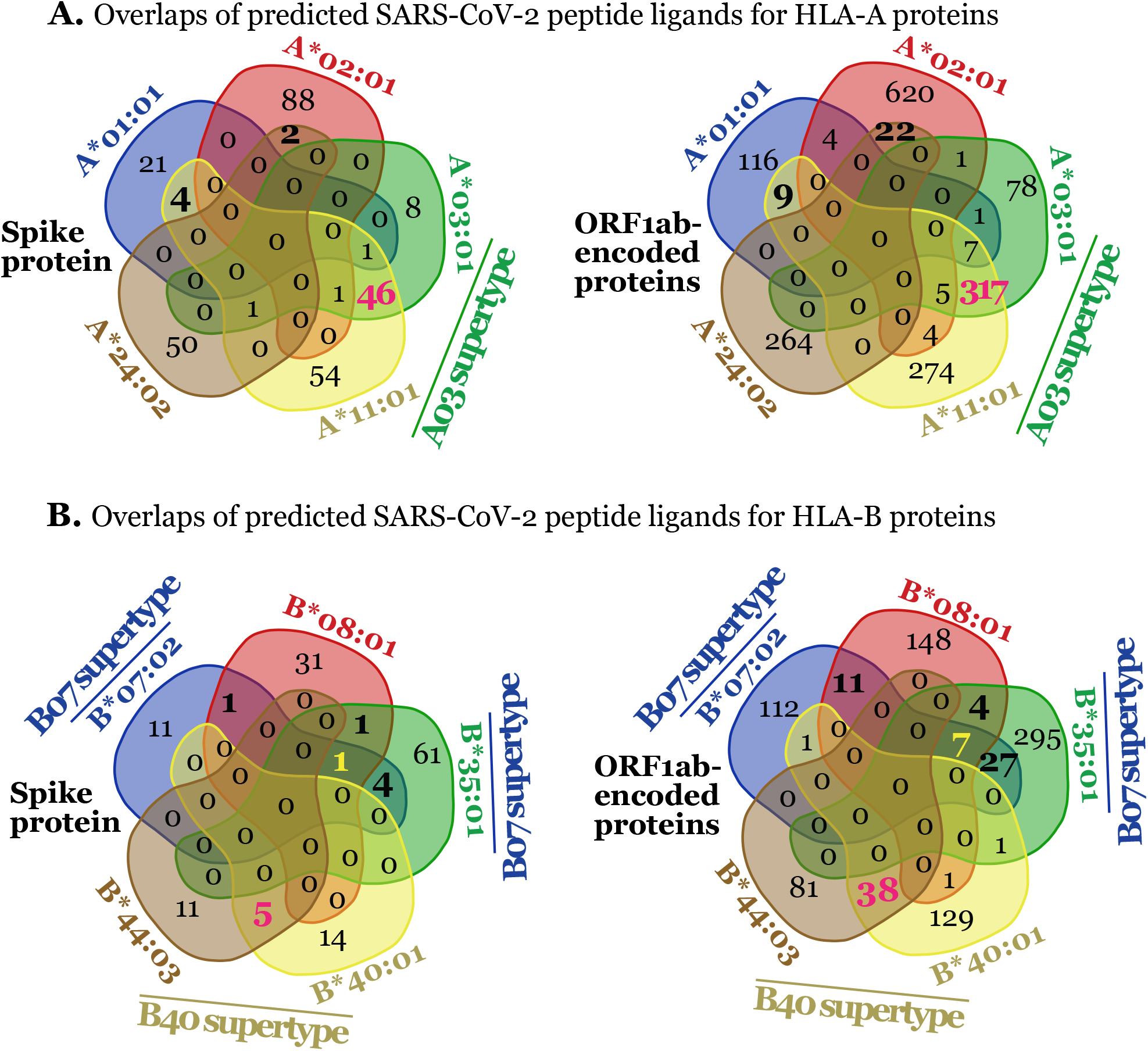
Dominant HLA class I proteins shared limited numbers of SARS-CoV-2 peptides. Numbers of predicted SARS-CoV-2 peptide ligands are shown for HLA-A proteins **(A)** and HLA-B proteins **(B)**.

Further, peptide-binding motifs of HLA class I proteins interact with anchor and preferred residues of peptide ligands to achieve high binding affinity (Fig. 3). The unique or shared anchor residues validates the binding of predicted SARS-CoV-2 peptides to specific HLA class I proteins or to HLA class I supertypes, respectively. Thus, the GibbsCluster program^53,54^ was applied to determine the residue patterns of predicted SARS-CoV-2 peptides. As a result, SARS-CoV-2 peptides showed conserved residues at positions 2 or 3 and 9, matching the reported anchor residues or preferred residues with corresponding HLA class I proteins (Fig. 3). HLA class I proteins assigned to the same supertypes shared similar anchor and preferred residues, as for supertypes B40 (B*40:01 and B*44:03), and to a less degree for B07 (B*35:01 and B*07:02)^55–60^. For A*03:01 with undefined anchor and preferred residues, this study showed a similar pattern of peptide binding to A*03:01 as to A*11:01, which display a sequence similarity of 96.15% (Fig. 1C) and belong to A03 supertype^52^. Therefore, predicted SARS-CoV-2 peptides form various patterns of anchor and preferred residues, which are shared within an HLA class I supertype (Fig. 3), as a unique conserved mechanism to implement HLA genotype-customized CD8^+^ T cell activation in various human populations.

**Fig. 3.**
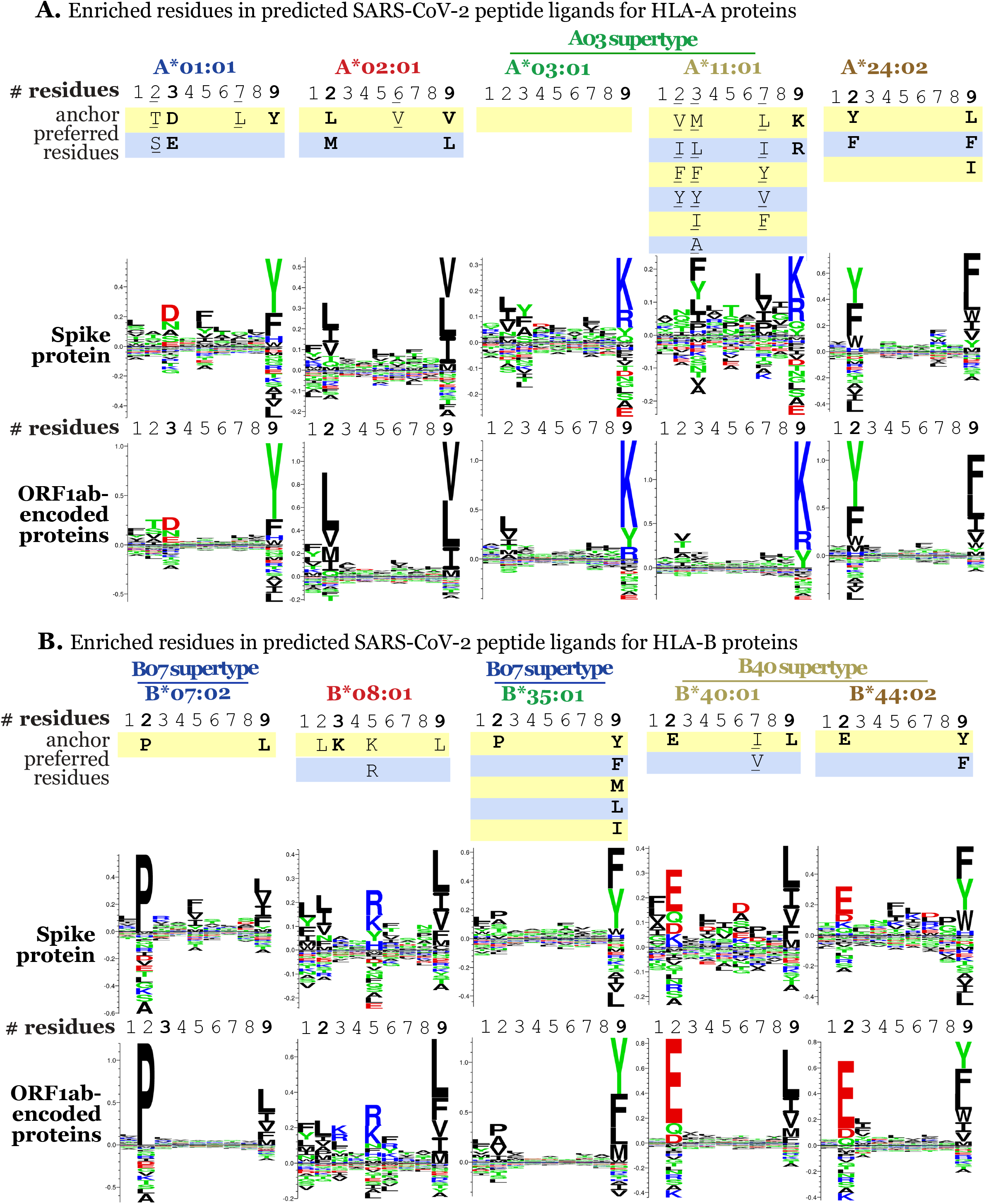
Enriched residues in predicted SARS-CoV-2 peptide ligands. Anchor and preferred residues for each HLA class I proteins are compared with enriched residues from predicted SARS-CoV-2 peptide ligands. Peptide ligands from the spike and ORF1ab-encoded proteins are shown for HLA-A proteins **(A)** and HLA-B proteins **(B)**.

### HLA class I proteins with weak peptide binding capacity incline to associate with severe infections

Multiple HLA class I proteins have been reported to associate with more severe diseases of viral infections. In SARS infections, B*46:01 was associated with severe cases in a study using a small sample size^44^ and the association of B*07:03 with severe SARS could not pass multiple tests^61^. In contrast, peptide binding to the HLA class I proteins of A03 supertype, including A*03:01 and A*11:01, was considered as potent stimulators for CD8^+^ T cells against SARS^45^, and the potential association of B*15:02 with resistance to SARS is not statistically significant in multiple tests^61^. Since individuals usually express two to four heterozygous or homozygous HLA class I A and B paralogous proteins, the potential association of one protein with severe diseases is likely balanced or interfered by other heterozygous HLA class I proteins in human population studies. The numbers of predicted peptide ligands in our study are supportive of a potential association of these HLA class I proteins with severity or protection of infections. For potent peptide binders potentially associated with resistance or protection, A*02:01, A*11:01, and B*15:03 sampled a large number of peptides (Fig. 4A). These results are consistent with a common stimulator of A*11:01 for anti-SARS CD8^+^ T cell responses^45^, a potential association of B*15:02 with the resistance to SARS^61^, and needs to further understand the effector phenotypes^27^ and kinetics^29^ of A*02-mediated CD8^+^ T cell responses in COVID-19. Belonging to the same supertypes (Fig. 4) and share high sequence homology (Fig. S2), A*02:02 sampled a high number of peptides as for A*02:01 (A02 supertype), and the similar was observed for B*15:01, B*15:02, and B*15:03, although a controversy remains in their assignment to one supertype or two supertypes^52,62^. Interestingly, SARS-CoV-2 peptides were predicted to be overlapped between A*02 and B*15 proteins or between A*11 and B*15 proteins (Fig. 4B), providing an additional possibility of identical peptides unusually shared by paralogous HLA class I proteins. On the other side, relative weak peptide binders B*07:03 and B*46:01 in this study were suggested potentially associated with severe SARS in some studies^61^ or with a small sample size^44^. B*07:02 and B*07:03 alleles (B07 supertype) sampled relatively fewer numbers of peptides in comparison to potent peptide binders such as A*02:02 or B*15:03, consistent with its difficulty to pass multiple tests in severe SARS^61^ if co-expressed with strong binders. Surprisingly, B*46:01 is predicted almost not binding to peptides from SARS-CoV-2 (Fig. 4A), suggesting a potential weak CD8^+^ T cell response against COVID-19 and consistent to its association with severe SARS infection^44^. A*25:01 binds a very limited number of SARS-CoV-2 peptides at low binding affinity, indicating a potential concern of weak responses in patients with an A*25:01 allele. These results allow us to predict and prepare to overcome weak peptide presenters in fighting human COVID-19 infections.

**Fig. 4.**
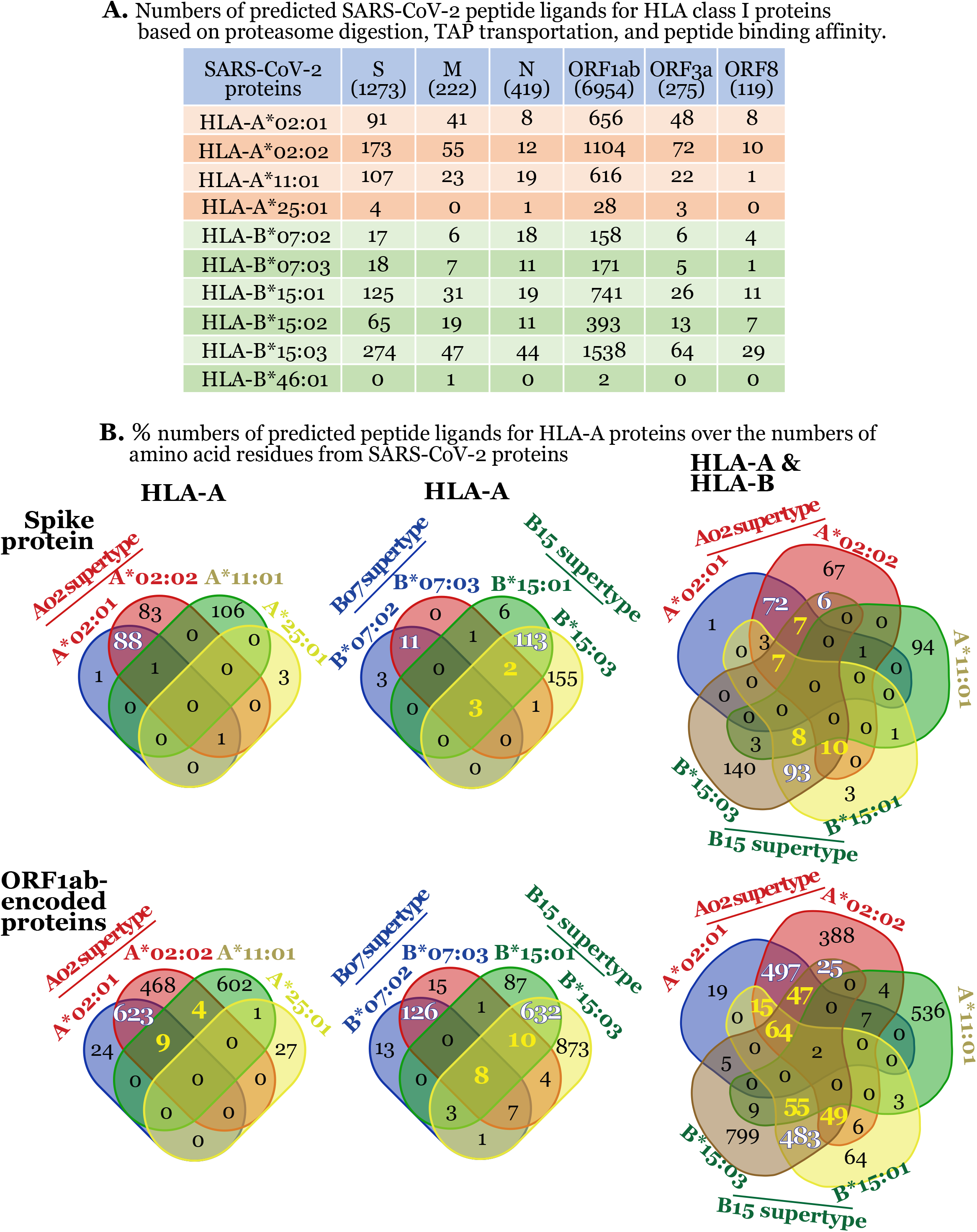
HLA class I proteins with weak peptide binding capacity inclined to associate with severe infections. Peptide ligands from immunogenic SARS-CoV-2 proteins were predicted for HLA class I proteins that were indicated relevant or irrelevant to the severity or protection of SARS. The numbers of predicted peptide ligands are listed for different dominant HLA class I proteins **(A)**. Venn diagrams show the numbers of predicted peptide ligands from the spike and ORF1ab-encoded proteins for HLA class I proteins **(B)**. White and yellow numbers are for overlaps by two and multiple HLA class I proteins, respectively.

### Overlapped HCoV and SARS-CoV-2 peptides suggests cross-reactive CD8^+^ T cells

Recent tests demonstrated that non-infected healthy donors contain CD8^+^ T cells responding to SARS-CoV-2 peptides^39,40^. Therefore, one can hypothesize that “common cold” coronavirus infection in humans can generate cross-reactive CD8^+^ T cells against SARS-CoV-2 infections. Moreover, cross-reactivity is more prominent and much stronger between SARS and SARS-CoV-2^40^. Our predicted numbers of overlapped peptides can overall explain this cross-reactivity. SARS and SARS-CoV-2 share various sequence similarity, such as 77.54% for S protein (Fig. S3) and 95.37% for ORF1b-encoded proteins (Fig. S4), contributing to the overlap of 69 peptides from spike protein and 438 peptides from ORF1b-encoded proteins between SARS with SARS-CoV-2 (Fig. 5). Since CD8^+^ T cells exposed to “common cold” coronaviruses are potentially reactive to SARS-CoV-2 peptides, the prediction of human coronavirus peptides contributing to this cross-reactivity will provide bases for cross-virus protection. However, HCoV-OC43 and SARS-CoV-2 share much lower sequence similarity at 33.06% for S protein (Fig. S3) and 62.32% for ORF1b-encoded proteins, contributing to 44 shared peptides (5%) from ORF1b-encoded proteins but nearly zero shared peptide from N, S, and ORF1a-encoded proteins between HCoV-OC43 and SARS-CoV-2 (Fig. 5). This limited number of cross-reactive peptides from HCoV is valuable for further functional investigations for cross protection.

**Fig. 5.**
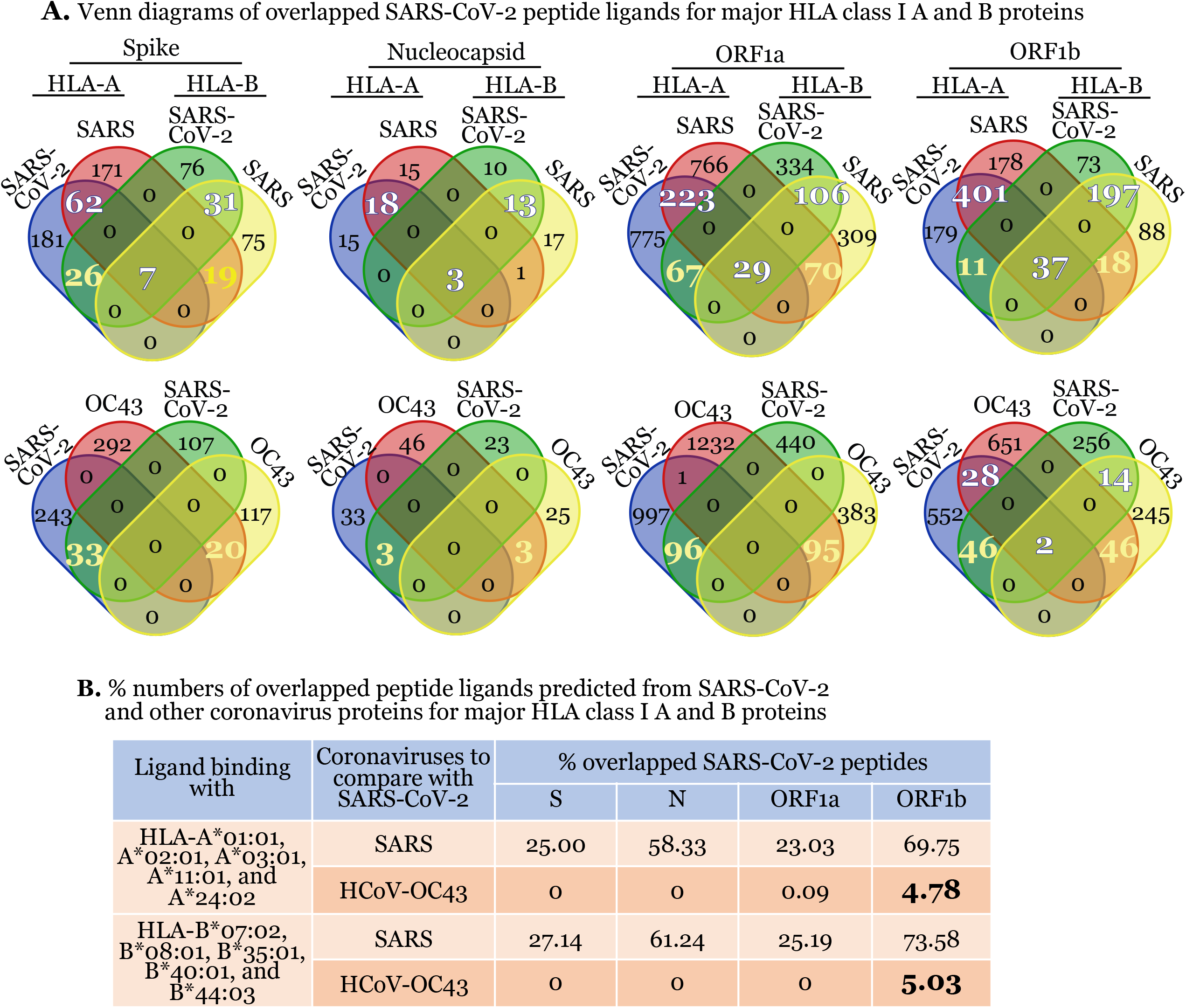
Overlapped peptides between SARS-CoV-2 and other coronaviruses. **A.** Venn diagrams show the numbers of identical predicted peptides shared between SARS-CoV-2 and other coronaviruses for HLA class I proteins. Large numbers show the overlaps between SARS-CoV-2 and other coronaviruses (white). Numbers of identical peptides loaded to both HLA-A and HLA-B proteins are shown in yellow. **B.** % of identical peptides between SARS-CoV-2 and other coronaviruses are listed. Bold % is for overlaps between SARS-CoV-2 and HCoV-OC43.

### Heterozygous HLA class I genotypes with high affinity peptides enhance CD8^+^ T cell responses in immune simulation

To predict whether high peptide binding affinity, peptide numbers, and HLA class I heterozygosity enhance CD8^+^ T cell responses, we use an agent-based simulator, C-ImmSim, to simulate antigen presentation for CD8^+^ T cell activation^63^. C-ImmSim mainly incorporated the impact of HLA-peptide binding, innate signaling, memory responses, and adjuvant effect in antigen presentation and CD8^+^ T cell activation, particularly differentiating the effect of heterozygosity or homozygosity of HLA genes^63^. First, we compared CD8^+^ T cell responses to one versus ten peptides with a high binding affinity (IC50<16nM) without involving innate or adjuvant factors. Results showed that one peptide is insufficient, but a combination of ten peptides, stimulated CD8^+^ T cell responses that became saturated in around two weeks (Fig. 6A). In this simulation, multi-epitopes stimulated more robust CD8^+^ T cell responses than one epitope, possibly due to very strong total binding avidity. Second, to more precisely test the avidity hypothesis, five peptides with the highest IC50 were compared with the other five peptides with the lowest IC50 as loaded to the same A*02:01 protein. Results further supported the conclusion that strong CD8^+^ T cell responses are more likely to be activated by multiple peptides with a high binding affinity (low IC50) to HLA class I proteins (Fig. 6B). Further, to determine whether heterozygous HLA-A or HLA-B proteins show an advantage to enhance CD8^+^ T cell responses than homozygous A*02:01 or B*44:02 alone, results showed peptides with similar or lower binding affinity stimulated stronger CD8^+^ T cell responses by heterozygous HLA-A or B proteins. Moreover, paralogous HLA-A and B proteins showed similar strength for CD8^+^ T cell activation to heterozygous HLA class I A or B proteins (Fig. 6C). Therefore, high peptide binding affinity, HLA gene heterogeneity, and paralogous HLA-A and B proteins provide advantages to stimulate a robust CD8^+^ T cell response.

**Fig. 6.**
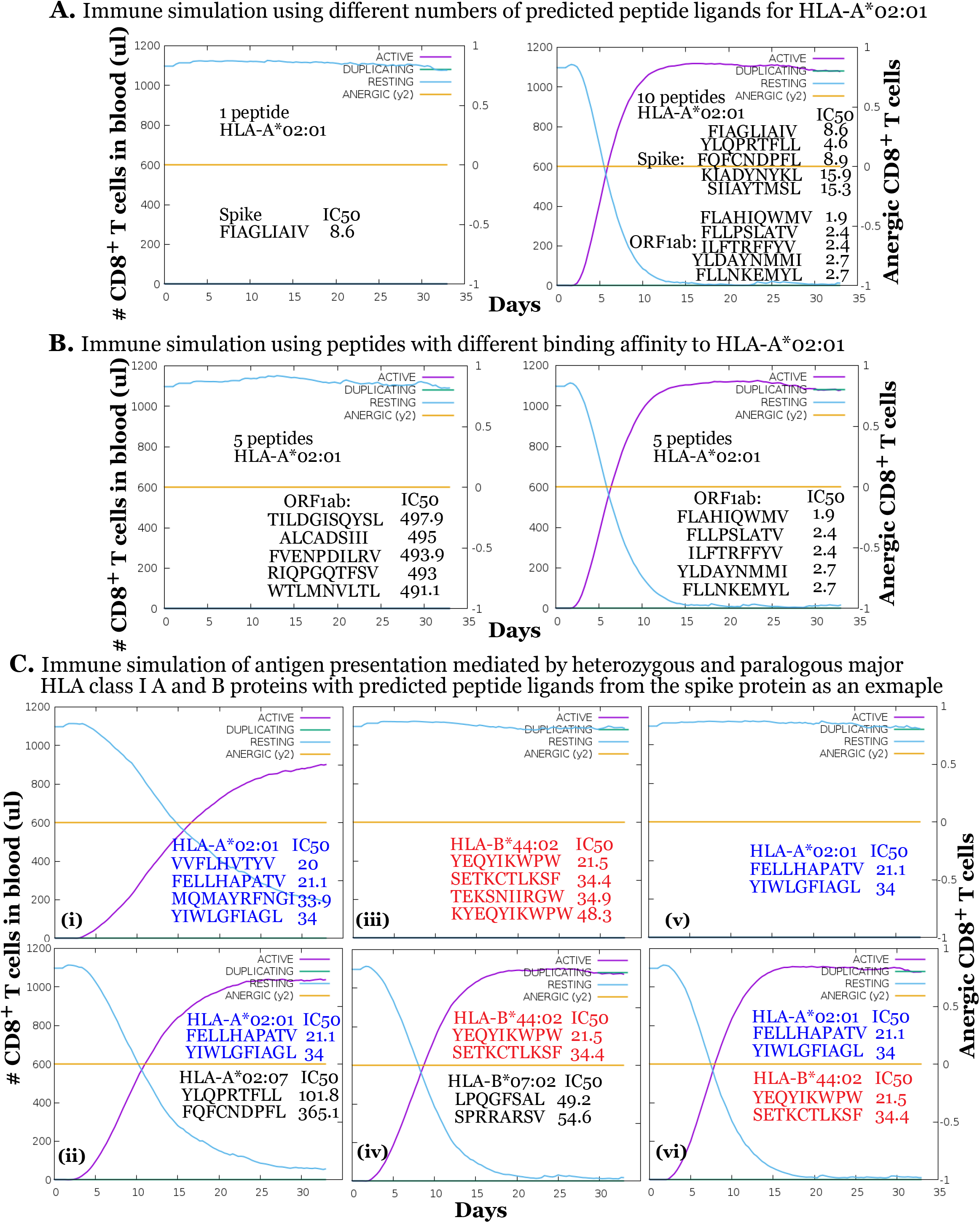
Heterozygous HLA class I genotypes with high affinity peptides enhance CD8^+^ T cell responses in immune simulation. Simulations of antigen presentation are compared with one and ten peptide ligands for A*02:01 protein **(A)**. Simulations of antigen presentation are compared between the high and low binding affinity of five peptide ligands for A*02:01 protein **(B)**. Simulations of antigen presentation are compared between homozygous **(i)** and heterozygous **(ii)** HLA-A proteins, homozygous **(iii)** and heterozygous **(iv)** HLA-B proteins, and with paralogous HLA-A and B proteins **(vi) (C)**.

### Heterozygous HLA class I genotypes with viral adjuvants rescue CD8^+^ T cell responses in immune simulation to weak peptide binders

Although B*46:01 and B*07:02 are potentially relevant to severe SARS^44,61^ and shown as weak peptide binders (Fig. 4), we speculated that CD8^+^ T cell responses in the individuals expressing one of these alleles can still be strongly stimulated by optimizing peptide immunization utilizing HLA class I heterogeneity and viral adjuvant. Strikingly, results showed that only three predicted peptides from multiple SARS-CoV-2 proteins bind to B*46:01 at a sufficient affinity (IC50<500nM) in comparison to hundreds or thousands of peptides sampled by other HLA class I proteins (Fig. 4A). One can imagine that an individual with a B*46:01 allele will be challenging for inducing CD8^+^ T cell activation, as this likely explains why B*46:01 allele is associated with severe SARS in a study using a small sample size^44^. Thus, it is critical to determine whether other HLA-A or B alleles in the same individuals facilitate overcoming this weakness. For example, HLA-A*02:07 and HLA-B*46:01 haplotype is present in around 3.34% of Asian pacific population^64^. Whether the peptides presented by HLA-A*02:07 can rescue or enhance CD8^+^ T cell responses mediated by the weak peptide binder B*46:01 is critical for inducing anti-COVID-19 immunity in this human population. By applying one additional peptide that is presented by a paralogous A*02:07 or by a heterozygous B*07:02 protein, results supported both strategies of antigen presentation overcame the disadvantage of B*46:01 protein for CD8^+^ T cell activation (Fig. 4A). Furthermore, the application of adjuvants such as an attenuated viral strain will help provide a cytokine environment mediated by innate cells and CD4^+^ T cells to facilitate the activation of peptide-specific CD8^+^ T cells (Fig. 7B and 7C). In this immune simulation, the viral adjuvant enhanced the proliferation of peptide-specific memory CD8^+^ T cells and balanced the production of IFNγ and inhibitory cytokines with extended kinetics. Therefore, a precise vaccination strategy considering potent peptide binders, heterozygous and paralogous advantages of HLA class I alleles, multi-epitopes of peptide antigens, and a viral adjuvant will induce strong anti-viral CD8^+^ T cell responses as tailored to individual and populational HLA class I genotypes.

**Fig. 7.**
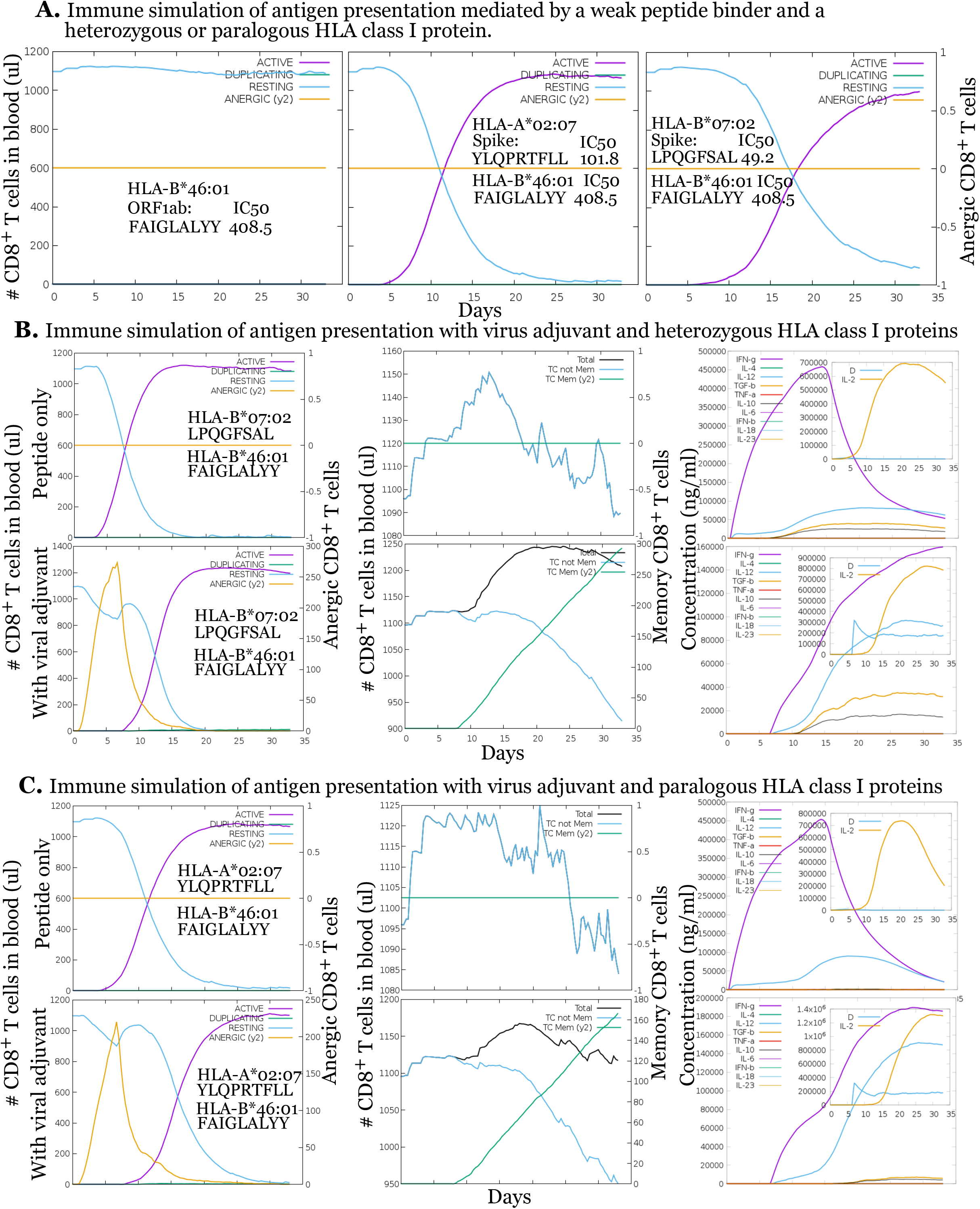
Heterozygous HLA class I genotypes with viral adjuvants rescue CD8^+^ T cell responses in immune simulation to weak peptide binders. Simulations of antigen presentation used B*46:01 only, mixed HLA-A and B*46:01, and heterozygous HLA-B alleles with B*46:01 **(A).** Simulations of antigen presentation used viral adjuvant and peptide ligands for both HLA-A and B*46:01 proteins **(B).** Simulations of antigen presentation used viral adjuvant and peptide ligands for heterozygous HLA-B proteins with B*46:01 **(C).**

## Discussion

More than two, three, and ten thousand HLA-A and B alleles have been detected in Asian, African, and European Americans, respectively^64^. The same viral proteins are expected to induce divergent CD8^+^ T cell responses in different individuals or populations, demanding a precise determination of efficacious peptide antigens presented by individual-specific HLA class I proteins. Remarkably extensive efforts are needed to structurally profile and functionally validate viral peptide pools for various HLA class I genotypes in ethnically diverse human populations. Computational prediction of these tens of thousands of peptides in terms of their binding affinity to each HLA class I proteins from tens of hundreds of HLA class I genotypes provides highly feasible alternatives to predict candidate peptides. Further, CD8^+^ T cell activation using these large numbers of combinations of predicted peptides and HLA class I genotypes can only be possibly performed using immune simulation to provide limited candidate strategies for further experimental and clinical tests. This study performed a computational prediction of peptide ligands and an immune simulation of CD8^+^ T cell activation using representative potent and weak peptide binders, including ten dominant HLA class I alleles and potentially SARS-associated HLA class I alleles. Results demonstrated that multi-epitope high-affinity peptides, tailored to heterozygous and paralogous HLA class I genotypes combined with a viral adjuvant, elicits strong CD8^+^ T cell responses, memory CD8^+^ T cell differentiation, and sustainable cytokine production.

Regardless of the degree of sequence diversity (Fig. S1C), key residues in peptide-binding motifs define the supertypes of HLA class I proteins^51,52^. HLA class I proteins of the same supertypes share peptide antigens and peptide-binding patterns, suggesting the conservation of anti-COVID-19 antigenic peptides in human populations. Briefly, large numbers of identical peptides overlapped between A*03:01 and A*11:01 (A03 supertype), B*44:03 and B*40:01 (B40) (Fig. 2), A*02:01 and A*02:02 (A02), B*07:02 and B*07:03 (B07), and B*15:01 and B*15:03 (B27) (Fig. In these HLA class I supertypes, A*02:01 and A*02:02 shared more than 50% of identical peptides, and three B*15 proteins shared large numbers of identical peptides (Fig. 4). Overlaps between paralogous HLA-A*02 and B*15 proteins also occurred at a much lower degree, implicating the presence of degenerate epitopes for CD8^+^ T cells^65^. Therefore, by defining HLA class I genotypes in targeted populations, similar or identical peptide antigens shared by HLA class I supertypes can be applied to vaccine design for those populations expressing HLA class I genes of the same supertypes.

Peptides with high affinity and avidity stimulate robust CD8^+^ T cell responses and aid weak peptide binder for T cell activation, as demonstrated by the immune simulation of antigen presentation in this study. One of our selected peptide, “FIAGLIAIV”, presented by HLA-A*02:01 has been validated for the activation of T cell from SARS patients^66^. Evidence was provided with insufficiency for stimulating CD8^+^ T cell responses using a single peptide (Fig. 6A) or five low-affinity peptides (Fig. 6B) in a vaccination setting in comparison to strong responses with ten peptides or five high-affinity peptides, respectively. Therefore, a high total peptide binding affinity (or avidity) is crucial for CD8^+^ T cell activation in vaccination.

Heterozygosity of HLA alleles is considered as a result of balancing selection to shape exceptional polymorphism of HLA proteins and increase host immunocompetence against highly diverse pathogens^67,68^. In this study, HLA class I heterozygosity was demonstrated to provide an advantage in presenting additional high-affinity peptides complementarily (Figs. 6 and 7). This heterozygous advantage allows the design of optimal peptide vaccination to achieve pronounced CD8^+^ T cell responses. For example, HLA-B*46:01 that was predicted to bind to two low-affinity peptides from ORF1ab-encoded proteins, in comparison to B*15:03 and B*07:02 predicted to bind 1538 and 158 peptides derived from the same SARS-CoV-2 proteins (Fig. 4), likely supports the theory of heterozygous advantage^68^, if these HLA class I alleles express in an individual. This study provided an example of immune simulation that CD8^+^ T cell responses in an individual expressing a weak peptide binder HLA-B*46:01 protein can be enhanced and rescued by the heterozygous expression of B*07:02 protein, which provided complementary capacity in peptide presentation to enhance CD8^+^ T cell activation (Fig. 7A). This heterozygous advantage has also been clinically demonstrated in individuals with heterozygous HLA class I loci A and B for protection against human T-lymphotropic virus type-1 (HTLV-1)^68,69^. Thus, HLA heterozygosity is an advantageous factor to be critically examined, particularly in the populations or individuals with weak CD8^+^ T cell responses to the virus- or protein-based vaccines, which can be rescued by providing a custom design of peptide antigens for the presentation of heterozygous HLA class I proteins.

Dual antigen presentation mediated by paralogous HLA-A and B proteins, as existing in all HLA class I haplotypes, further enhances CD8^+^ T cell responses. This dual antigen presentation elicits more robust CD8^+^ T cell responses than the antigen presentation mediated by a single HLA-A or B protein for a similar number of peptides (Fig. 6C). Using the weak peptide binder B*46:01 as an example, peptides presented by HLA-A proteins can be applied to rescue the weak CD8^+^ T cell responses mediated by HLA-B*46:01 protein. For example, in clinical trials of anti-viral vaccination, 3.34% of the Asian Pacific population co-expressing HLA-A*02:07 and the weak peptide binder HLA-B*46:01^64^ can be considered to have additional peptide-based vaccination to elicit optimal CD8^+^ T cell responses, as customized to this specific HLA class I genotype (Fig. 7). Moreover, viral co-vaccination or adjuvant can be applied to boost more robust or stabilized CD8^+^ T cell responses for the haplotype containing B*46:01 in human populations, together with antigen presentation mediated by either heterozygous or paralogous HLA class I A and B proteins (Fig. 7).

In summary, precision vaccination strategies targeting CD8^+^ T cells can be customized on HLA class I genotypes and peptide affinity for human populations to circumvent weak peptide binders in T cell activation. Dependent on HLA class I genotypes from an individual or a population, high-affinity peptides can be matched to heterozygous or paralogous HLA class I A and B alleles together with adjuvant as customized vaccination strategies. This key-to-lock model^33,34^ explains why vaccine efficacy results are usually very different between animal experiments and human trials, because antigen-presenting molecules and peptide antigens for T cell activation are entirely different between animals, including primates, and humans. However, clinical trials to determine the efficacy of a vaccine is time- and labor-consuming, and economically challenging, due to the requirement for double-blind design and the observation of the protection under natural infection conditions in a long-term process. Computational simulation of antigen presentation, using this study as an example, provides valuable candidate strategies for HLA genotype-customized vaccine design using known viral protein sequences. It is reasonable to speculate that the antigen presentation tailored to HLA genotypes of patients is advantageous to overcome weak CD8^+^ T cell reactivity and facilitate achieving a high efficacy of protection in anti-SARS-CoV-2 vaccine trials clinically.

## Materials and Methods

1. **Protein sequences were accessed** from GenBank with accession numbers of QHU79204.1 (spike or S protein), QLI50116.1 (membrane or M protein), QHU79211.1 (nucleocapsid or N protein), QJQ39966 (open reading frame 1ab or ORF1ab), QJQ27841 (ORF1a), QJQ39969 (ORF3a), QJQ27861 (ORF8), Protein sequences of human beta coronavirus OC43 isolate include AGT51680.1 (S protein), AIX10719.1 (N protein), YP_009555238.1 (ORF1ab), and QBP84755.1 (ORF1a). Protein sequences of SARS include AAP51227.1 (S protein), AAP51234.1 (N protein), QJQ39966 (ORF1ab), QJQ27841 (ORF1a). ORF1b fragment is defined by the non-overlapped portions of amino acid sequences between ORF1ab and ORF1a. HLA class I protein sequences for peptide binding prediction were accessed from the website of The European Molecular Biology Laboratory (EMBL)-European Bioinformatics Institute (EBI) (https://www.ebi.ac.uk/ipd/imgt/hla/allele.html).
2. **The alignment of protein sequences** was performed using a hierarchical clustering approach (http://multalin.toulouse.inra.fr/multalin/)^50^. Neutral consensus residues are labeled in red, low consensus residues are labeled in blue, and high consensus portions are labeled in grey as a background. Upon sequence alignment, percentages of identical sequences are calculated and listed.
3. **Affinity-based prediction of peptide ligands for HLA class I proteins.** The model NetMHCpan was chosen because this model has been widely used to predict the binding affinity of peptide sequences with 8-11 residues to various HLA class I proteins. NetMHCpan calculates a 50% maximal inhibitory concentration (IC50) based on the maximal concentration of test peptides to compete off a probe peptide for binding to the same HLA class I protein^46^. NetMHCpan prediction has been performed accurately in recent antigen peptide prediction to integrate mass spectrometry-eluted ligands and binding affinity of peptides^70,71^. NetMHCpan is also available for all MHC alleles to analyze dominant and disease-associated HLA class I alleles. This study used the NetMHCpan 4.0 program, which is a default method in the website of Immune Epitope Database (IEDB.org) for predicting peptide ligand binding to HLA class I proteins. This prediction was not limited to the most typical length of peptide sequence (9 amino acids) but expanded to cover a possible length of 8 to 11 residues.
4. **Proteasome degradation** mainly considers the degradation mediated by immune proteasomes, which are induced by Interferon γ to generate antigenic peptides. Immune proteasomes degrade SARS-CoV-2 proteins to polypeptides, which have a high potential to bind to MHC Class I molecules. The prediction was based on *in vitro* proteasomal digestion of enolase and casein proteins as described^48^ to determine the numbers of potential cutting sites at the C-termini induced by various proteasome proteases. In this prediction, we used the Netchop v3.0 “C-term” model to remove any peptides that were not predicted via proteasomal cleavage of the peptide’s C terminus^72,73^. A conserved threshold of five was used to select peptide ligands with five or more cutting sites in proteasome degradation to generate peptides for further canonical MHC class I antigen processing.
5. **Transportation by TAP transporters.** Upon proteasome digestion, cytosol peptides are required to be transported by peptide transporters across the endoplasmic reticulum membrane to be loaded to the heavy chain of HLA class I proteins. It has been shown that a high affinity of TAP transporter binding to a peptide translates into high transport rates^49^. Notably, both proteasome and TAP predictions were developed using experimental data for HLA proteins and were highly suitable for predicting viral peptide ligands for HLA class I proteins. Interestingly, TAP ligands with high binding affinity have an increased chance of being cleaved by the proteasome, and TAP specificity has evolved to fit the digestion specificity of proteasomes^73^. Thus, the prediction of TAP transporter binding to peptides links the processes of proteasome digestion and peptide loading to HLA class I proteins. A TAP score estimates −log(IC50) values (log=base 10) of peptides for binding to TAP. The cutoff of IC50 at 1 nM was used to remove peptides with higher IC50 or lower TAP binding affinity^49^.
6. **Alignment of peptide residues.** We align short peptides using the computational program Gibbs (http://www.cbs.dtu.dk/services/GibbsCluster/), which performs two essential tasks simultaneously, alignment and clustering^54^. We used Gibbs to de-convolute binding motifs of SARS-CoV-2 peptide datasets. GibbsCluster simultaneously clusters and aligns peptide data as a powerful tool for unsupervised motif discovery. Results return optimal clusters with characterized sequence motifs for each cluster based on multiple parameters, including adjustable penalties for small clusters, adjustable penalties for overlapping groups, and a trash cluster to remove outliers. The results of cluster peptides are shown with the frequency of specific amino acids. The sequence motifs derived from the best solution are displayed in the form of sequence logos generated with Seq2Logo. Colors highlight different categories of amino acids, including acidic (red), basic (blue), uncharged (green), and other residues (black). The pre-identified anchor and preferred residues of previously defined peptide ligands for HLA class I proteins were obtained from (http://www.syfpeithi.de/bin/MHCServer.dll/FindYourMotif.htm) to compare with the enriched clusters of predicted SARS-CoV-2 peptides.
7. **Venn diagram** was constructed using the website (http://bioinformatics.psb.ugent.be/cgi-bin/liste/Venn/calculate_venn.htpl) for four-way and five-way Venn diagrams. Sequentially identical or overlapped peptide sequences are defined as identical or overlapped peptides.
8. **Immunity simulation** was performed using C-ImmSim program (http://150.146.2.1/C-IMMSIM/index.php?page=0)^63^. This prediction program implements a Celada-Seiden model as a logical description of the mechanisms making up humoral and cellular immune responses to a genetic antigen by incorporating known principal factors. These core factors include MHC restriction, clonal selection by antigen affinity, antigen presentation in cytosolic or endocytic pathways, cell-cell interactions, T cell differentiation, and T cell memory for T cell responses. Simultaneously, C-ImmSim considers the helper function of CD4^+^ T cell to antibody production and CD8^+^ T cell responses. C-ImmSim has been used to predict immune response in HIV infections, which have been validated with published clinical trials involving antiretroviral therapies^74^. This study utilized C-ImmSim to predict whether peptide binding affinity, peptide numbers, and combined HLA class I alleles impacted CD8^+^ T cell response to SARS-CoV-2 peptides. Input factors included peptide sequences, HLA class I alleles, incorporation of a viral vaccination as an adjuvant for this study. Results are shown with the curve of CD8^+^ T cell proliferation and cytokine production to predict potential effector T cell responses and memory T cell differentiation stimulated by HLA class I-restricted peptides.

## Acknowledgement

Thanks to the American Lung Association for funding support (IA-629987).

## Author Contributions

S.H. developed research workflow, analyzed data, and wrote the manuscript. M.T. provided inputs on viral vaccination and manuscript writing.

## Supplementary Information

**Fig. S1.**
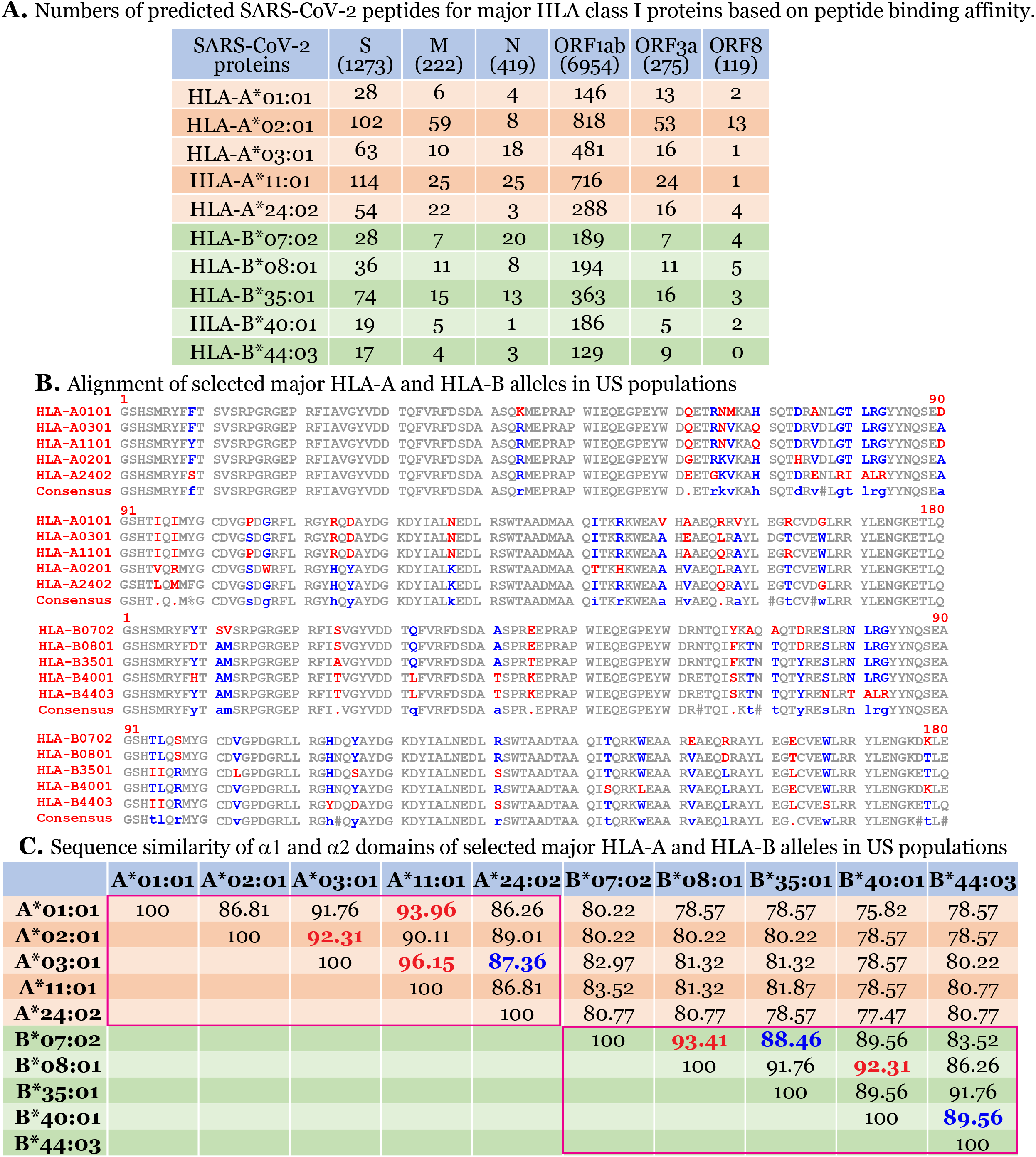
Predicted SARS-CoV-2 peptide ligands only based on binding affinity to HLA class I proteins and sequence similarity in HLA class I proteins. Numbers show predicted peptide ligands from immunogenic SARS-CoV-2 proteins only based on binding affinity to major HLA class I proteins **(A)**. Alignment of α1 and α2 domains of these HLA-A and HLA-B proteins was performed using Clustal Omega program with residues colored in grey (identical residues), red (highly diverse), and blue (less diverse) **(B)**. % identical residues are listed (C).

**Fig. S2.**
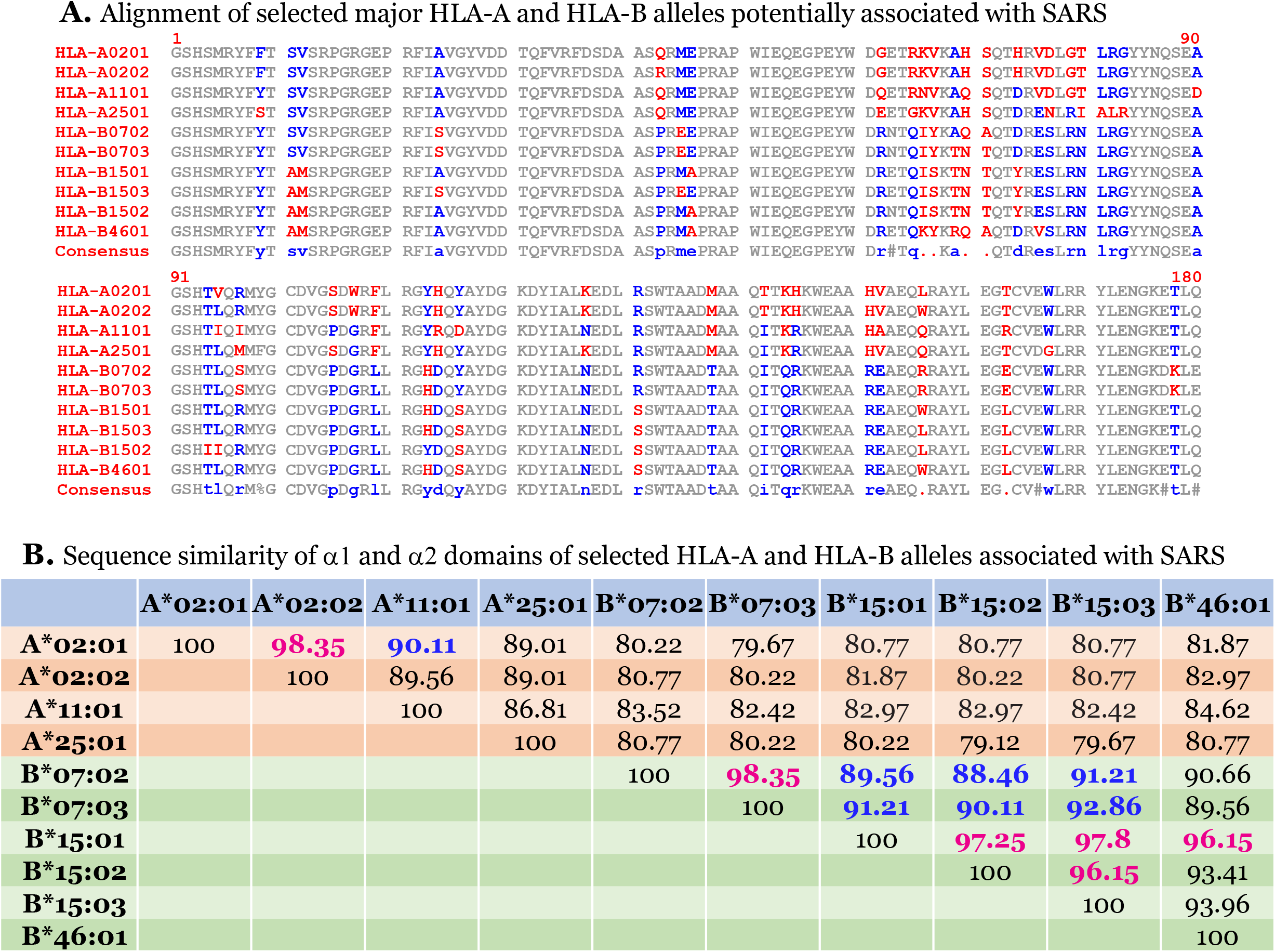
Alignment and sequence identity of HLA class I proteins relevant or irrelevant to SARS infections. **A.** alignment of α1 and α2 domains for HLA class I proteins that were indicated relevant or irrelevant to the severity or protection of SARS. Residues are colored in grey (identical residues), red (highly diverse), and blue (less diverse). **B.** % identical residues are listed.

**Fig. S3.**
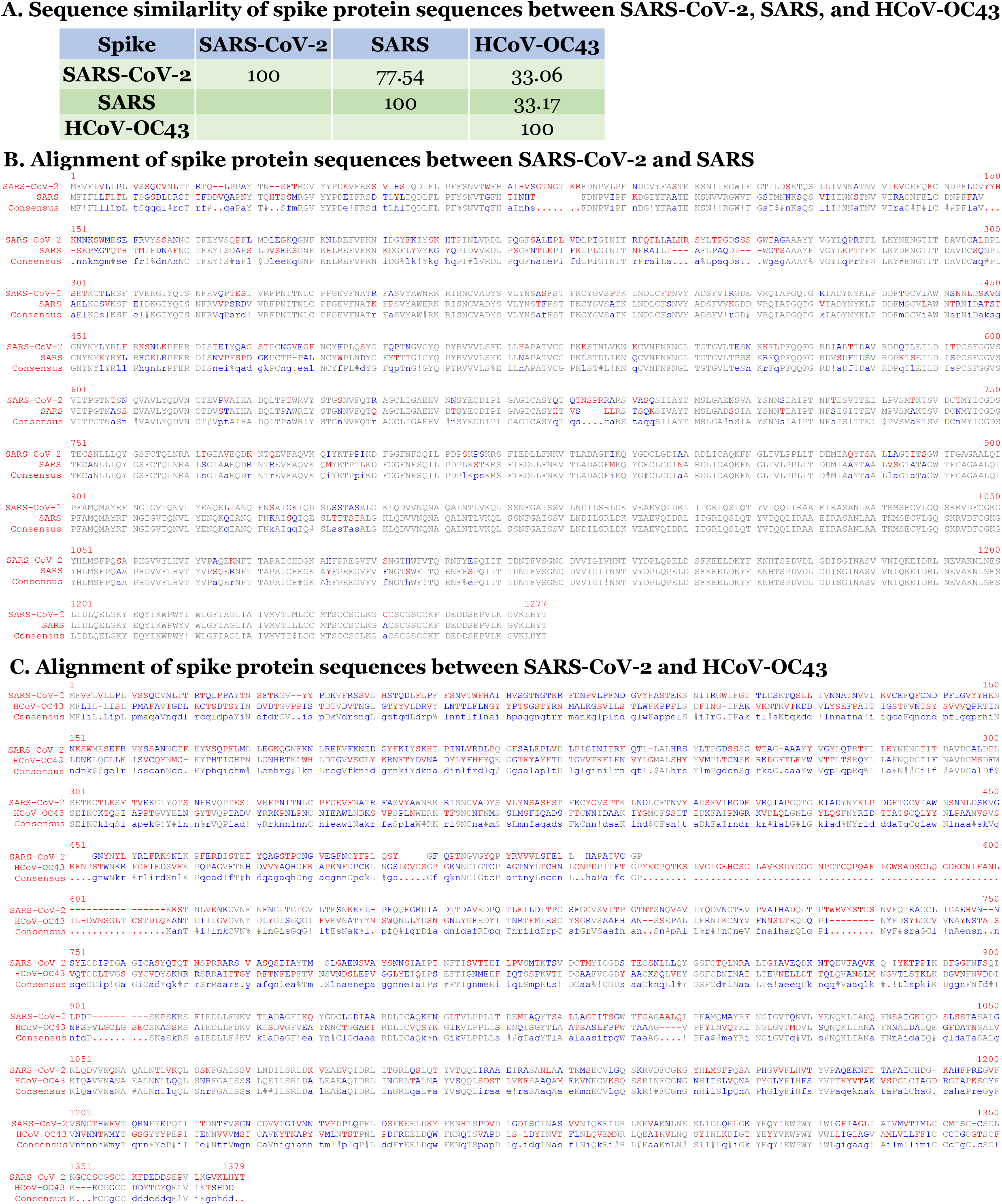
Alignment and sequence identity of spike protein between SARS-CoV-2 and other coronavirus. **A.** % sequence identity is listed. **B.** alignment of S protein between SARS-CoV-2 and SARS. **C.** alignment of S protein between SARS-CoV-2 and HCoV-OC43.

**Fig. S4.**
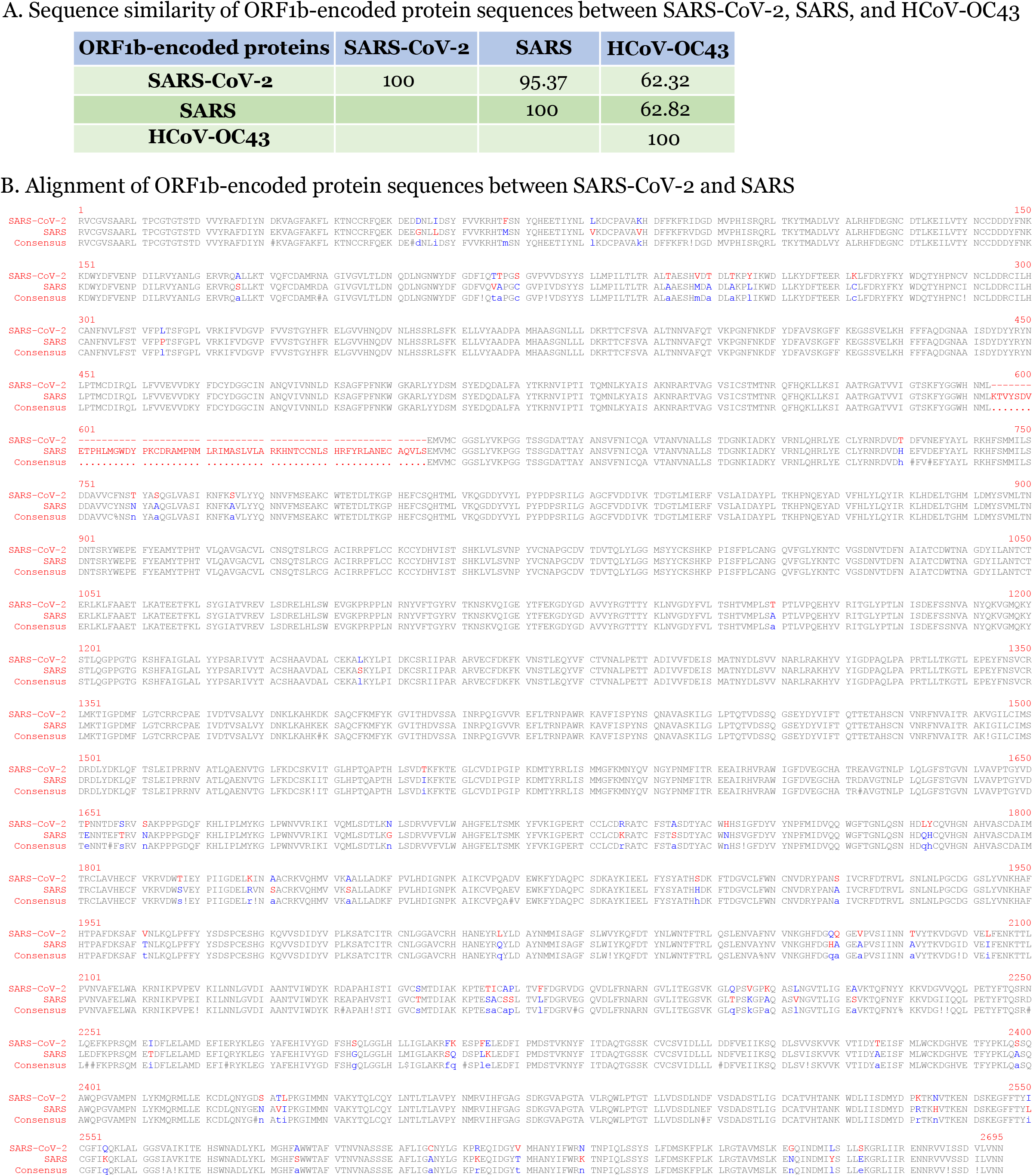
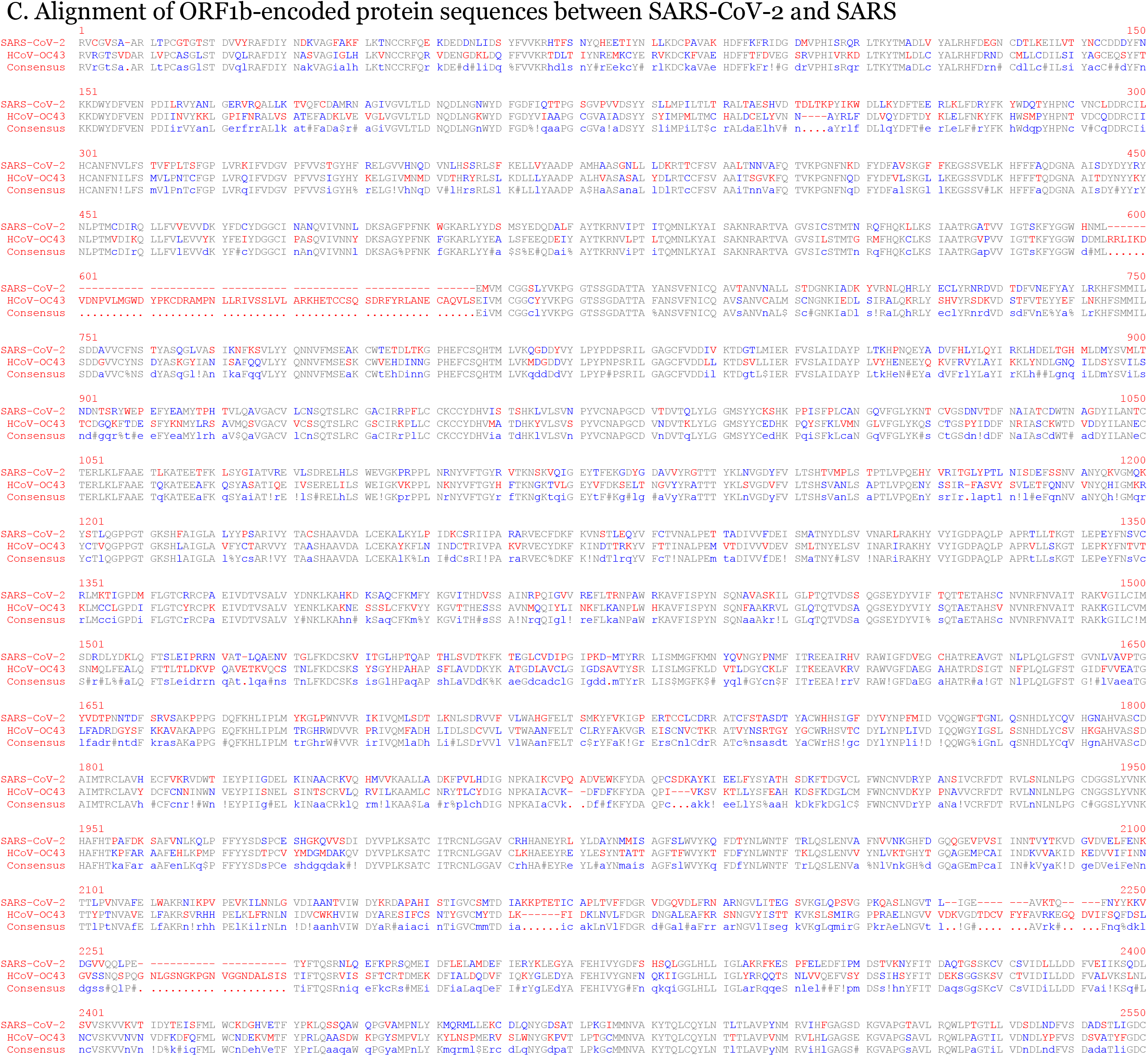
Alignment and sequence identity of ORF1b-endoced proteins between SARS-CoV-2 and other coronaviruses. **A.** % sequence identity is listed. **B.** alignment of ORF1b-endoced proteins between SARS-CoV-2 and SARS. **C.** alignment of ORF1b-endoced proteins between SARS-CoV-2 and HCoV-OC43.

## Notes

### Competing Interest Statement

The authors have declared no competing interest.

